# Taxonomic Profiling and Populational Patterns of Bacterial Bile Salt Hydrolase (BSH) Genes on Worldwide Human Gut Microbiome

**DOI:** 10.1101/260794

**Authors:** Ziwei Song, Yuanyuan Cai, Xue Wang, Xiaoxuan Lin, Yingyun Cui, Jing Shang, Liang Jin, Jing Li

## Abstract

Bile salt hydrolase (BSH) in gut bacteria can hydrolyze conjugated bile salts to unconjugated bile acids and amino acids. Thereby play a crucially important role in host health by reducing serum cholesterol levels, preserving bile acids balance and regulating various metabolism as signaling molecules. Here we present the taxonomic identification of BSHs in human microbiota and elucidate the abundance and activity differences of various bacterial BSHs among 11 different populations. For the first time, we have revealed BSH are distributed in 154 intestinal bacterial strains within 33 genera in human microbiota. However, these BSHs present obviously differentiation for the sequence identity being from 28.6% to 100%, and the 32.7% bacteria strains having more than one paralogs of BSHs with dissimilarity. Therefore, we reclassified the BSHs from the different genera into 6 phylotypes basing on their phylogenetic tree, and demonstrate the significant abundance patterns of BSH phylotypes among different populations. From the enzyme activity comparison, the representative sequence of BSH-T3 was shown highest enzyme activity in 6 phylotypes. Meanwhile, BSH-T3 sequences which all distributed in *Lactobacillus* show highest abundance in Chinese and Austrian. The information illustrated by this study is useful for investigating the population differences of bile acid metabolism related diseases, and further giving a new suggestion on selection of probiotics or development of pharmaceutical proteins based upon the activity of BSH phylotypes to regulate host metabolism and maintain fitness.

## Introduction

Bile acids are well known for regulating cholesterol balance, and the disorder of bile acids enterohepatic circulation could cause gallbladder (Wang et al., 2008) or gastrointestinal diseases (Copaci et al., 2005). It is also pointed that bile acids metabolism is associated with diabetes (Staels and Kuipers, 2007), obesity (Joyce et al., 2014) and cardiovascular diseases (Porez et al., 2012). Bile acids are synthesized from cholesterol in the hepatocytes, which are further conjugated with the amino acids glycine or taurine to bile salts. It is noteworthy that the amphiphilic combination of bile salts is essential to absorb fat in the intestine. However, excessive bile salts are toxic to intestinal bacteria (Alvarez-Sola et al., 2017).

Bile salt hydrolase (BSH) in gut microbiome could catalyze the hydrolysis of bile salts into deconjugated bile acids for preserving the balance of bile acids metabolism. Moreover, the deconjugated bile acids are serving as signaling molecules facilitates the secretion of GLP-1 (Parker et al., 2012), activating multiple receptors (Degirolamo et al., 2014; S., 2008; Studer et al., 2012), and affecting a variety of metabolic processes and diseases (Li and Chiang, 2014).

BSH have already been identified in some microbial genera, including *Lactobacillus* (Corzo and Gilliland, 1999), *Bifidobacterium* (Kim et al., 2004), *Enterococcus* (De Filippo et al., 2010), and *Clostridium sp p*(Coleman and Hudson, 1995), *etc*. Interestingly, it is reported that two BSH enzymes (BSH1 and BSH2) from *Lactobacillus salivarius LMG14476* are striking different in positive cooperatively, catalytic efficiency and substrate preference (Bi et al., 2013). Furthermore, there is reported 3 distinct BSH have been identified in the same strain, i.e. *L.johnsonii PF01* (Chae et al., 2013). These studies indicate that the paralogs number of BSH genes in some strains can be variable.

Given the important role of gut microbiota in the bile acids metabolism, it is vital to investigate which bacteria having BSH genes and how the abundance and activity variations of BSH within human gut microbiota systematically. Due to the rapid developing of next generation sequencing technology, the public databases, such as

The Human Microbiome Project (HMP) (Human Microbiome Project, 2012), META genomics of the Human Intestinal Tract (MetaHIT) (Li et al., 2014), providing the baselines and variances of human gut metagenomic data. From previous studies, different populations exhibit considerable gut microbiota variances of due to different genetic backgrounds, environments, and especially huge difference in dietary habits (Liu et al., 2017; Liu et al., 2016), which also helpful to investigate the variation patterns of BSH in human gut microbiota.

In this study we used computational and biological approaches to investigate the taxonomic, populational and functional patterns of BSHs in human gut microbiota with the following underlying aims: (1) to identify which bacteria holding BSHs in human gut microbiota; (2) to investigate how the sequential, structural, and biological activity variations of the BSHs from different bacteria; (3) to explore what factors might affect the patterns of BSHs in human microbiota.

## Results

### Sequence and structure comparisons of classical BSHs

Firstly, we obtained a total of 144 BSHs from NCBI which were distributed in 7 genera, including *Lactobacillus*, *Bifidobacterium*, *Listeria*, *Enterococcus*, *Carnobacterium*, *Eggerthella* and *Eubacterium* (Figure S2A). However, only 10 BSH 3D structures were reported in PDB. To probe the difference of BSHs in different genera, we compared the sequences and structures of BSHs from three classic genera, *Bifidobacterium*, *Lactobacillus* and *Enterococcus*.

From the result of sequences comparison, the number of amino acids in BSHs from *Bifidobacterium* was 315, from *Lactobacillus* was 317 and from *Enterococcus* was 322 (Figure 1A). The average sequences identity of these BSH sequences was 54.64%.

**Figure 1.**
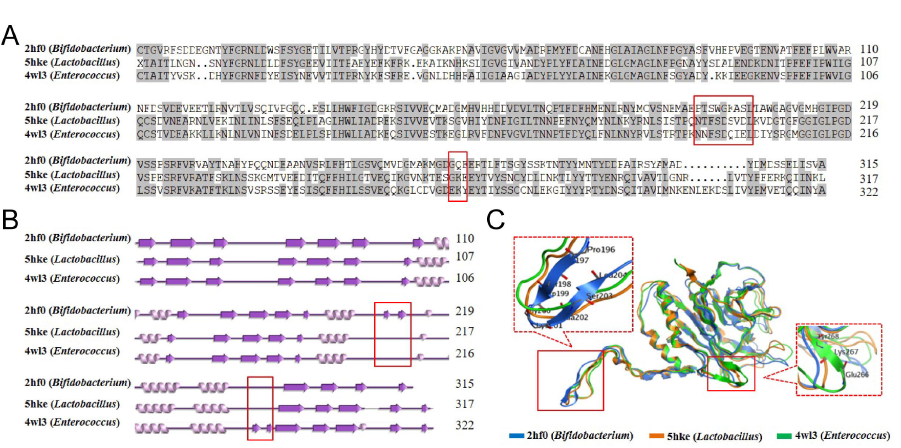
Sequence and structure comparison of three representative BSHs from different bacterial genus. (A) Multiple sequences alignment of BSHs, Grey background means BSHs identity ≥50%. (B) Secondary structure comparison of BSHs. (C) 3D structure comparison of BSHs. The red frames are shown the major different parts in these three BSHs.

The second structures showed that all these BSHs contain 5 α-helixes. The only difference was the number and position of β-sheets (Figure 1B). Detailedly, BSHs from *Bifidobacterium* and *Enterococcus* contain 19 β-sheets, while BSHs from *Lactobacillus* only keep 17 β-sheets. Two special β-sheets of BSH in *Bifidobacterium* was from Pro196 to Leu204, and in *Enterococcus* was from Glu266 to Tyr268.

The 3D structure comparison of these BSHs was similar as secondary structure comparison, there were two specific β-sheets on BSHs from *Bifidobacterium* and *Enterococcus*, respectively (Figure 1C). It was reported that the activity sites of BSHs were near Asn175 and Arg228 (Chand et al., 2017), which was not close to the structural differential sites. The results indicated that the relationship of BSH structure and enzyme activity should be investigated in following study.

### Taxonomic identification and NP investigation of BSHs

To explore the difference of all BSHs from bacteria in human microbiota, 210 reference BSHs were constructed from HMP database. All these BSHs were assigned to 33 genera from 4 Phyla, *Actinobacteria*, *Euryarchaeota*, *Firmicutes* and *Proteobacteria* (Figure 2A). In addition, 75% of BSHs could be assigned to the genus of *Lactobacillus*, *Enterococcus*, *Lister*, *Bifidobacterium* and *Bacillus*. Particularly, the

**Figure 2.**
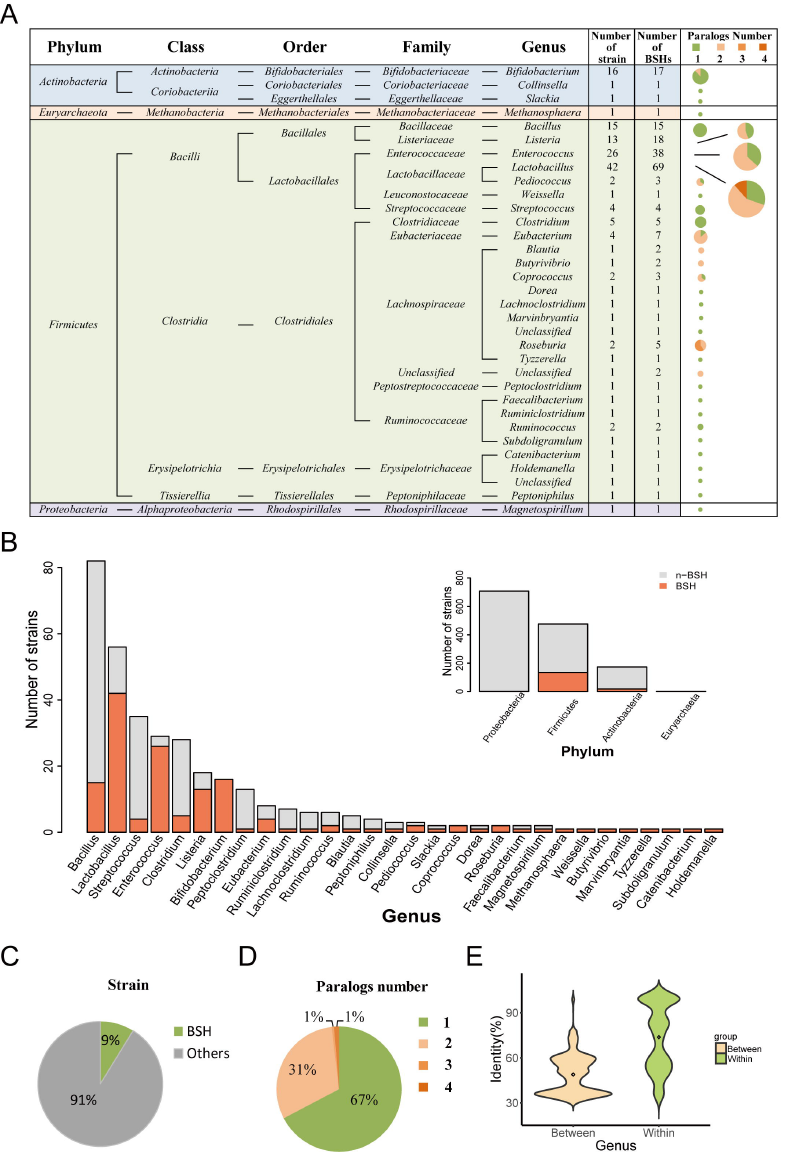
Taxonomic characterization of BSHs obtained from the HMP. (A) Taxonomic identification of BSHs and NP in different genus. (B) Quantity and proportion of BSHs containing bacteria stains in phylum and genus level. (C) The proportion of strains that have BSHs in all strains from HMP database. (D) The proportion of different paralogs number in all BSHs from HMP database. (E) Density distribution of BSHs between genus and within genus in HMP database.

*Lactobacillus* contained 69 of 210 BSHs. Furthermore, these 210 reference BSHs were distributed in only 154 strains due to the existent of paralogs (Figure 2A). There were totally 11 genera of bacteria have paralogs. Especially, one strain of *Roseburia* contained 3 BSHs, and two strains of *Lactobacillus* contained 4 BSHs (Table S1).

The number and proportion of bacteria stains that containing BSHs in phylum and genus were calculated, respectively. Most BSHs containing bacteria were belonging to *Firmicutes*. In particular, all strains of *Bifidobacterium* contained BSHs, and the BSHs containing bacteria stains certainly occupy a large proportion in *Lactobacillus*, *Enterococcus* and *Lister* (Figure 2B). It was shown that total 9% of bacteria strains in HMP microbiota reference genome contained BSHs (Figure 2C).

The proportions of different NP were shown that most of strains (67.31%) contained only one BSHs, 30.77% of strains have two BSHs, 0.64% have three and only two strains of *Lactobacillus* have four BSHs (Figure 2D).

The identity of BSH between genera demonstrated mainly distributed in two intervals, one interval was about 35%, and the other was about 60%. But there was also a lower identity of BSHs within genus gathered at about 50%. These results indicated that distinguish BSHs by genera might not be a rational method due to the paralogs of BSHs in harboring bacterial genome (Figure 2E).

### Populational comparison and multivariable adjusted analyses of BSHs

We further checked the distribution and abundance of BSHs. There were 95 out of 210 reference BSHs found in human gut metagenome based on sequence similarity searching. The relative abundance of these 95 reference BSHs were calculated (Table S3).

Considering the influence of individual condition, we compared the difference of BSHs distribution among age, gender and body mass index (BMI) groups by using single factor analysis. The relative abundance of BSHs was significant higher (*p*=0.024) in female gut microbiota than in male (Figure 3A). The correlation analysis was shown that there was no relevance between the relative abundance of BSH and age (Figure 3B) or BMI (Figure 3C) of individuals.

**Figure 3.**
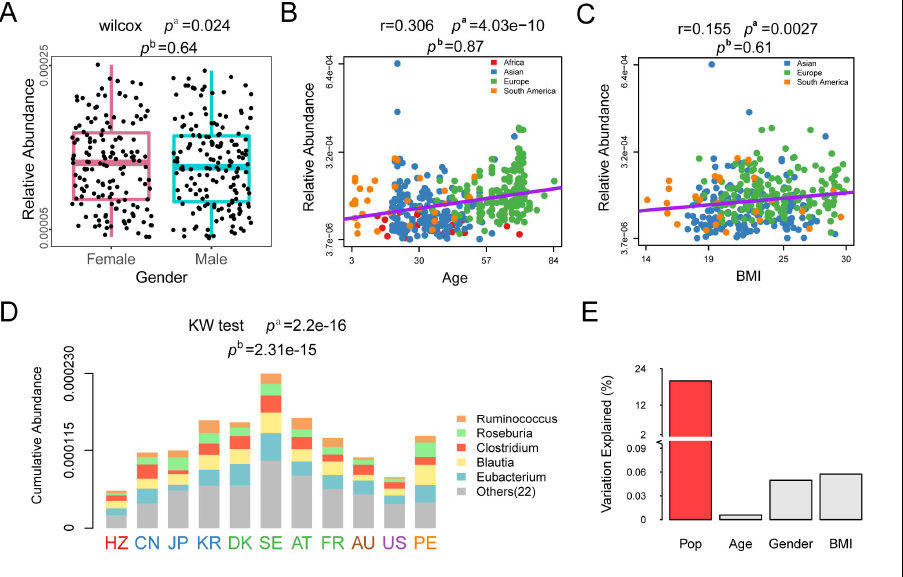
Effects of multiple factors on relative abundance of BSHs. (A) Relative abundance of BSHs in different gender in 9 populations. (B) Correlation between BSHs relative abundance and age in 8 populations. (C) Correlation between BSHs relative abundance and BMI in 7 populations. (D) The cumulative abundance and genera distribution of BSHs in different populations. (E) The multiple regression model of BSHs cumulative abundance against individual condition including 4 factors. *p*^a^ values were calculated by nonparametric univariate method (Mann-Whitney-Wilcoxon test) corrected for false discovery rate. *P*^b^ values were calculated by multivariable adjusted analyses using linear regression model for one factor data with remaining 3 factors data.

From the cumulative abundance distribution of BSHs in different populations (Figure 3D, Table S3), the SE population exhibited the highest BSHs cumulative abundance (2.30×10^−4^), and HZ population exhibited the lowest BSHs cumulative abundance (5.52×10^−5^). It was probably due to HZ population behaved a more simpler composition of gut microbiota than other populations, as HZ people were the last remaining populations in Africa that live a hunter-gatherer lifestyle. On the continent level, cumulative abundance of BSHs in Europe (DK, SE, AT, FR) populations shown higher level, i.e. 1.34×10^−4^ ~ 2.30×10^−4^ comparing the average abundance of 1.31×10^−4^. Moreover, it was shown that the majority of BSHs are identified from 5 genera, i.e. *Eubacterium*, *Blautia*, *Clostridium*, *Roseburia* and *Ruminococcus* in the stack columns, which represent about 69.1% total abundance of BSHs in human gut metagenomes.

Multivariable regression analysis was implemented on abundance of BSHs against population, age, BMI and gender. Firstly, the effect of gender, age and BMI on BSHs abundance were not significant after adjusted (Figure 3A, 3B, 3C). Then, it was shown that the significant association was only between the BSHs cumulative abundance and populations (adjusted *p*=2.3e-15, Figure 3E), populational structure could explain the highest variations of total abundance variances of BSHs among 621 healthy samples. These results were consistent with the report that the disparity of gut microbiota composition in different populations mainly caused by geography.

### Reclassification and variation patterns of BSHs

Given the BSHs within genera shown broad range of sequence dissimilarity due to the NP of BSHs in many strains, the BSHs abundance pattern at genus level might not reflect the functional variations and phenotype association well. Thus, we reclassified the 95 BSHs in 11 human gut microbiome into 6 phylotypes based on the phylogenetic tree (left panel of Figure 4, Figure S3). It was shown that there are 4 major phylotypes, i.e. BSH-T1 to BSH-T4. Meanwhile, BSH-T01 and BSH-T02 were two clusters that couldn’t be classified by hierarchical algorithm but relative close in the phylogenetic tree.

**Figure 4.**
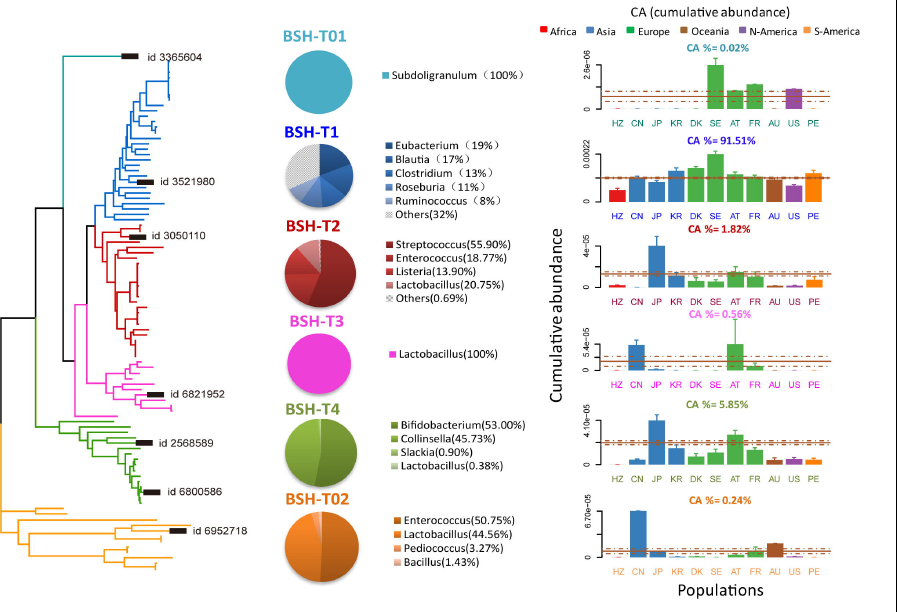
Taxonomic characterization and populational patterns of reclassify BSHs. Left panel is the phylogenetic tree of BSHs in 11 populations, different colors represent 6 reclassify BSH phylotypes. The highlight sequences with id are selected for enzyme activity experiment. The pies in the middle panel are shown the proportions of genera in different BSH type. Bar plots in the right panel indicate the cumulative abundance of BSH phylotypes in different populations.

Among the genera distribution (Figure 4, middle panel) and population patterns (Figure 4, right panel, Table S4) of each phylotype, BSH-T01 and BSH-T02 keep only 0.26% cumulative abundance (CA) of BSHs, which were mainly distributed in *Enterococcus* and *Lactobacillus*. BSH-T1 contained 91.51% CA of BSHs were mainly distributed in *Eubacterium*, *Blautia*, *Clostridium*, *Roseburia* and *Ruminococcus*, and could be found in the gut microbiota of all 11 populations. BSH-T2 contained 1.82% CA of BSHs, and the major genera of this phylotype were *Streptococcus* and *Lactobacillus*. It was noteworthy that this phylotype cannot be found in gut microbiota of CN population. Sequences of BSH-T3 were all from *Lactobacillus*, which was shown higher cumulative abundance in CN population and AT population. BSH-T4 included 5.85% CA of BSHs, which were mainly distributed in *Bifidobacterium* and *Collinsella*. The cumulative abundance of BSH-T4 in gut microbiota of JP population was shown particularly higher than other populations.

Above all, the BSHs from *Enterococcus* and *Lactobacillus* were shown special phylotype distribution, i.e. it could be found in BSH-T2, BSH-T3, BSH-T4, and BSH-T02. This result has corresponded with that most of strains from these two genera harboring more than one paralogs of BSHs (Figure 2A). Thus, different paralogs of BSHs within genus could have sequence dissimilarity, which might lead to variable functional roles of these two genera in bile acid metabolism.

### Molecular docking and enzyme activity comparisons in 6 BSH phylotypes

To compare hydrolysis capacity among 6 types of BSH, BSHs behaved the highest relative abundance in each phylotypes were chose to do molecular docking and enzyme activity assay (Table S5). As lower binding energy (BD Energy) of molecular docking indicated more stable of the complex. The stability of the complex was BSH-T3 > BSH-T2 > BSH-T4 > BSH-T02 > BSH-T01 > BSH-T1 (Figure 5A). The active sites of BSH-T3 docked with bile acids were Tyr185 and His214, 5 residues from 4 chains formed 6 hydrogen Bond with GDCA and 5 hydrogen Bond with TDCA. The most hydrogen bonds lead to the lowest BD energy BSH-T3 shown than other BSHs. So that, BSH-T01, which was formed only 4 hydrogen Bond with GDCA and 1 with TDCA, shown the highest BD energy (Figure 5A, Figure S4).

**Figure 5.**
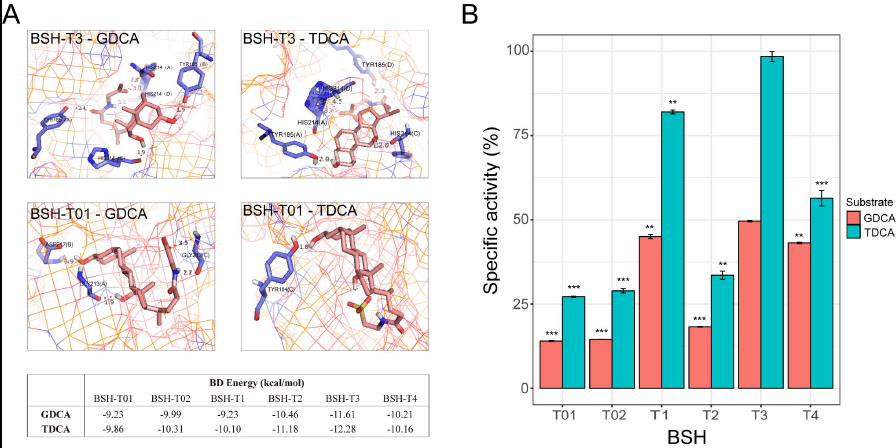
Molecular docking and enzyme activity comparation. (A) The Autodock predicted binding models of BSH-T3 and BSH-T01 with bile acids (GDCA, TDCA). Binding energy predicted by Autodock are displayed at the bottom. (B) The specific activity (%) comparison of different BSH type. Using the highest enzyme activity as 100%. * *p* < 0.05, ** *p* < 0.01, *** *p* < 0.05 versus BSH-T3.

Purified BSH of 6 phylotype were used to compare the enzyme activity. A single protein band indicating all BSH were obtained after SDS-PAGE analysis of purified protein samples (Figure S4). Results of BSH activity comparison were BSH-T3 > BSH-T1 > BSH-T4 > BSH-T2 > BSH-T02 > BSH-T01 (Figure 5). In both virtual docking and biological enzyme activity assay, BSH-T3 was shown the highest activity, which BSHs were all distributed in *Lactobacillus*. Comparatively, the BSH-T01 and BSH-T02, i.e. the two type including broad bacteria without good classification, were shown lower enzyme activity consistently in computational and in vitro assessment. It noteworthy that the BSH-T1, the type with the highest abundance of BSHs in human gut microbiota, exhibited the secondary highest enzyme activity in vitro assay but not in the docking test. Though the biological assay should more likely reflect the reality of enzyme activity, this conclusion should be verified in the following in vivo assay.

## Discussion

In our study, BSH protein sequences were used to do BLASTP for their sensitivity in sequences searching compared with gene sequences. The average of sequence identity of BSHs between genera was 49.7% and within genus was 66.92% (Figure S2B). Thus, the BLOSUM45 matrix and identity 45% as cutoff was used for building the reference BSHs to make sure all BSHs could be screened from HMP database. Meanwhile, the BLOSUM62 with parameters of e-value 1e-5 and identity 62% as cutoff were used for identifying BSHs sequences in each individual to the accurate taxonomic assignments of BSHs in human gut microbiota.

From our results, the taxonomic characteristic of reference BSHs was shown that 34% of the BSHs contained bacteria behaved paralogs. The presence of paralogs may cause BSH protein sequences and functional differences in the same bacteria strain. Thus, we reclassified the 95 BSHs to 6 phylotypes based on the phylogenetic tree, and the taxonomic characterize of different phylotypes were diverse. Interestingly, 4 of 6 phylotypes all have BSHs from *Lactobacillus*, which was coordinated with the result of *Lactobacillus* behaved the most NP in 33 genera containing BSHs.

BSH-T3, all identified from *Lactobacillus*, showed the highest enzyme activity in both virtual and in vitro assay. Among the 3 typical BSH with available 3D structures (Figure 1), the BSH (PDB id: 5hke) from *Lactobacillus* are also belonging to BSH-T3. Comparing to other types of BSHs, there are lacking four β-sheets beside the active sites (Tyr185 and His214) for BSH-T3. This structural character might lead to the His214 of BSH-T3 easily establishing the stable hydrogen bond with GDCA or TDCA, which also illustrate the higher affinity of BSH-T3 to the substrates. Moreover, the highest bioactivity of BSH-T3 was also provided an evidence for the beneficial effect of *Lactobacillus* products, i.e. The abundance of *Lactobacillus* exhibits the positive association with the increasing ability of bile acid metabolism.

We have analyzed multiple factors correlations of BSHs abundance in gut microbiota of worldwide healthy individuals (Figure 3). After adjusted, BSHs abundance was no significant difference between populations of different gender (*p*=0.64), and no significant correlation with age (*p*=0.87) or BMI (*p*=0.67), it was exhibited significant difference only in different populations (*p*=2.31×10^−15^). These results gave an important baseline of the BSH variations among healthy individuals. The different diet, climate and local habitats, etc. with different lifestyle of worldwide populations were led to the huge populational variances of gut microbiota. Therefore, although not directly related, we still constructed the association between the populational BSHs abundance with WHO released phenotype, which might shed light on high incidence of bile acid metabolism related phenotypes in some particular populations (Figure S5). However, all BSHs abundance only shown a weak correlation with the death rate of diabetes (r=-0.55, *p*=0.079), only BSH-T02 abundance shown significant correlation with mean blood cholesterol (r=-0.72, *p*=0.0042) and death rate of CVD (r=0.87, *p*=0.0053) among 6 phylotype. Besides the limitation of two unrelated datasets, it might indicate that the relationship between BSHs abundance and bile acid metabolism related phenotypes were indirect and intricate, which need to explore in the following studies.

In this work, we describe the distribution of bacteria that behaved BSHs, discuss distribution characteristic of BSH abundance in worldwide populations. After the above series of analysis, we propose a new method to reclassify the BSHs, and compare the activities between different types. However, future work is required to understand the subtle interplay between microbe-generated BSH and bile acid metabolism related health or disease in the host.

## Author Contributions

JL and LJ conceived and designed the study. ZS, YC, XW, XL, YC, JS, LJ and JL collected data and performed the analyses. JL and ZS interpreted the data and wrote the paper. All authors have read and approved the final version of the manuscript.

## Acknowledgements

J.L. acknowledges financial support from the National Natural Science Foundation of China (Grant No. 31670495) and a Project Funded by the Priority Academic Program Development of Jiangsu Higher Education Institutions (PAPD) and Top-notch Academic Programs Project of Jiangsu Higher Education Institutions (TAPP). The funders had no role in study design, data collection and analysis, decision to publish, or preparation of the manuscript.

J.S. acknowledges financial support from the One Hundred Person Project of The Chinese Academy of Sciences, Applied Basic Research Programs of Qinghai Province (No. Y229461211), Science and Technology Plan Projects in Qinghai Province (No.2015-ZJ-733), Science and Technology Plan Projects in Xinjiang (No. 2014AB043), Prospective joint research project of Jiangsu Province (BY2016-078-02) and Xinjiang science and technology assistance program(2016E02067).

## Method and Materials

### Sequences and structures comparison

The protein sequences and 3D structures of BSHs from *Bifidobacterium* (PDB ID: 2HF0), *Lactobacillus* (PDB ID: 5HKE) and *Enterococcus* (PDB ID: 4WL3) were downloaded from the protein data bank (PDB, http://www.rcsb.org/pdb/home/home.do). The secondary structures of these BSHs were downloaded from PDBsum (http://www.biochem.ucl.ac.uk/bsm/pdbsum). Multiple amino acid sequence alignments of the BSHs were carried out using the DNAMAN (v 8.0). Comparison of 3D structures was carried out using MOE (v 2014).

### Taxonomic characterization

The initial 144 query BSH protein sequences were collected from NCBI (https://www.ncbi.nlm.nih.gov/) with the keyword “bile salt hydrolase” and BLASTP (v2.2.29+)(Camacho et al., 2009) according to parameters of e-value=1e-5 was used to make their own alignments to obtain the identity interval of the BSHs between genera and within genera. The reference genomes of 1,751 bacterial strains covering 1,253 species were obtained from HMP in September 2014. The genes and related proteins from these bacteria genomes were predicted by MetaGeneMark (v2.8)(Zhu et al., 2010), and the taxonomic information of these genes and proteins was directly extracted from the strain names. The initial 144 BSHs are BLASTP with parameters of e-value=1e-5 and 45% sequence identity as cutoff was used for searching all BSHs in HMP database.

### Public available metagenomic sequence data

The metagenomic sequence data of individuals were collected from Hadza (HZ)(Rampelli et al., 2015; Schnorr et al., 2014), China (CN)(Qin et al., 2012; Qin et al., 2014), Japan (JP)(Nishijima et al., 2016), South Korea (KR)(Lim et al., 2014), Denmark (DK)(Le Chatelier et al., 2013), Sweden (SE)(Karlsson et al., 2013), Austria (AT)(Feng et al., 2015), France (FR)(Zeller et al., 2014), the USA (US) (HMP DACC, http://www.hmpdacc.org), Peru (PE)(Obregon-Tito et al., 2015), and Australia (AU)(PRJEB6092). To construct metagenomic datasets of healthy individuals from each country, the data for individuals with BMI ≥30, those designated with the following diseases based on the literature: inflammatory bowel disease, type 2 diabetes, or colorectal cancer, and infants <3 years old were excluded from the data. Although we could not access the metadata for individuals from KR, AU and US, we used all data from individuals with an average BMI of 24 ± 4 for this cohort. Finally, a total of 621 healthy individuals from the 11 countries were selected for analysis (Table S2).

### Metagenomic analysis based on different populations

All the raw sequencing reads were assessed and filtered using FASTX-Toolkit, and the high quality microbiome sequencing reads were assembled by the package SOAPdenovo2 (v2.04)(Ruibang L, 2012). After assembling, the contigs with at least 500bp were further used to predict the genes by MetaGeneMark (v2.8), and then a non-redundant protein set was constructed by pair-wise comparison of all protein sequences within populations using BLASTP (v. 35x1)(Kent, 2002) with 95% identity and 90% overlapping thresholds. The relative abundance of each protein sequence in each individual was calculated by the number of read pairs mapped to the gene over the length of protein sequence and divided by the summary of sequence abundance per individual(Qin et al., 2012). Cumulative abundance (CA) was calculated by the sum of the same individuals and the same genera using the relative abundance divided by the number of people in each population. The BSHs were identified in above population database using BLASTP with parameters of e-value=1e-5 and 62% sequence identity and 210 query sequences were identified.

### Phylogenetic tree

The phylogenetic tree was built by using Maximum likelihood (ML) method in MEGA software (v 7.0). Dendroscope (v 3.4.7) was used to embellish the phylogenetic tree, adjust labels and fill colors as needed.

### Related physiological/diseases data

The mean blood cholesterol (2008), obese BMI (2014), deaths rate (per 100000 data) of diabetes (2012) and deaths rate (per 100000 data) of cardiovascular diseases (2012) by country were downloaded from World Health Organization (WHO, http://www.who.int/en/). The average of data was used to do the correlation analysis when the data is sexually differentiated.

### Homology modeling and Molecular docking

The online software, Protein Homology/analogY Recognition Engine V 2.04 (Phyre2, http://www.sbg.bio.ic.ac.uk/phyre2) was used to predict the homologous structure of BSHs using the intensive mode (Kelley et al., 2015). Ligands including GDCA (C_26_H_43_NO_5_) and TDCA (C_26_H_45_NO_6_S·2H_2_O) were obtained from the ZINC database (http://zinc.docking.org/) and for docking analysis. The Autodock program (version 4.2.6) (Morris et al., 1998) was employed to generate an ensemble of docked conformations for each ligand bound to its target. BSH was bond with substrate by tetramers. We used the genetic algorithm (GA) for conformations search, and performed 100 individual GA runs to generate 100 docked conformations for each ligand.

### Materials, bacterial strains and vectors

Glycine deoxycholic acid (GDCA), taurodeoxycholic acid (TDCA), glycine and taurine standards were obtained from Solaibao (Beijing, China). pET28a (+) was used for expression of His-tagged (6x) recombinant BSH (Genscript, Nanjing, China). The resulting fragments containing the BSH genes was ligated into the NcoI/XhoI-digested vectors pET28a (+), resulting in 6 recombinant plasmid pET28a (+) - *bsh* transformed into *E. coli* BL21 (DE3) competent cells.

### Expression of BSH genes

The BSH-recombinant bacteria were inoculated into Luria-Bertani (LB) broth which containing 50μg•ml^−1^ kanamycin and incubated at 37 °C until the optical density at 600 nm (OD_600_) reached about 0.5. Expression of the BSH genes was induced by the addition of isopropyl β-D-thiogalactoside (IPTG, 0.5 mmol•l^−1^), then incubated at 37 °C (BSH-T5) for 4 hours or 16 °C (BSH-T01, BSH-T02, BSH-T1, BSH-T2, BSH-T3, BSH-T4) for 16 hours. Cells are harvested by centrifugation at 8000 rpm for 10 min at 4 °C. Then the cell pellet was resuspended (1: 10) in suspension (0.05M Tris, 0.05M NaCl, 0.5 mM EDTA, 5% Glycerol) and disrupted by sonication (alternating pulses: on for 3 s, off for 3 s; 60% amplitude). At last, supernatant fluid was collected to purify after harvested (8000 rpm, 10 min, 4 °C).

### Purification of the BSH

The disrupted liquid was harvested at 8000 rpm for 15 min at 4 °C. This BSH soluble fraction was placed in a nickel-nitrilotriacetic acid (Ni^2+^-NTA) agarose column (GE Healthcare, USA). The column was equilibrated with 20 mmol•l^−1^ imidazole. The present and purification of BSHs were confirmed by SDS–PAGE.

### BSH Assay

BSH specific activity was determined by measuring the release of amino acid from conjugated bile salts. The amounts of amino acids liberated by BSHs are determined by the ninhydrin assay (Dong et al., 2015). Protein concentrations were determined with the Bradford Protein Assay kit (Beyotime, Nanjing, China), and bovine serum albumin was used as standard. The purified BSHs lyophilized powders were proper diluted by 0.1M PBS (pH 6.0). We used 10 μl of BSH liquid and 10 μl of bile acids (200mM) and 180 μl of reaction buffer (0.1 M PBS, pH 6.0), and 10 μl of liquid paraffin were added. After the reaction was incubated at 37 °C for 30 min, 200 μl of 15% (w/v) trichloroacetic acid (TCA) was added to terminate the reaction, and the sample was centrifuged to remove the precipitates. Then, 10 μl of the supernatant was completely mixed with 190 μl of ninhydrin reagent, and boiled for 15 min. The tubes were cooled to room temperature and the absorbance at 570 nm was measured. A standard curve was prepared with glycine or taurine. One unit of BSH activity was defined as the amount of enzyme that released 1 mol of amino acids from the substrate per min.

### Statistical analysis

All values were expressed as mean ± SD. Statistically significant differences between groups were determined by the wilcoxon rank-sum test. The correlation analysis was done with the Spearman correlations test. Multiple comparisons were performed by multiple regression analysis. All the statistical analyzes in this paper were performed using R (v 3.3.2), *P* < 0.05 was considered as statistically significant.

**Figure S1.**
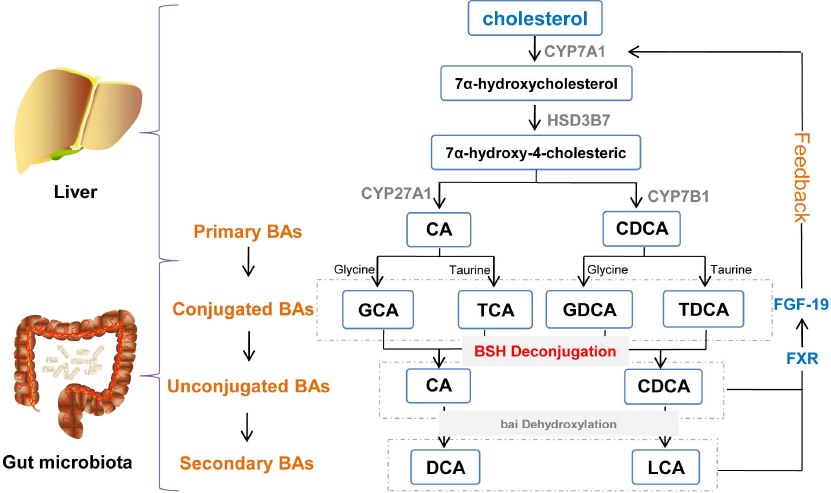
Related to Figure 1. The enterohepatic circulation of bile acids. In the liver, cholesterol is converted to 7α-hydroxycholesterol by CYP7A1. The 3β-hydroxysteroid dehydrogenase (HSD3B7) converts 7a-hydroxycholesterol to 7a-hydroxy-4-cholesteric, which eventually is converted to primary bile acids (BAs) by CYP7B1 and CYP27A1. The primary BAs (CA and CDCA) usually conjugated with the glycine or taurine to conjugated Bas (GCA, TCA, GDCA and TDCA), and transfer to intestine. BSH in gut microbiota would catalyze the hydrolysis of conjugated BAs into deconjugated BAs. Then, microbial bai dehydroxylase removes a hydroxyl group from C-7 and converts CA to DCA and CDCA to LCA. These secondary Bas could active the feedback pathway by FXR, thus to maintain BAs balance.

**Figure S2.**
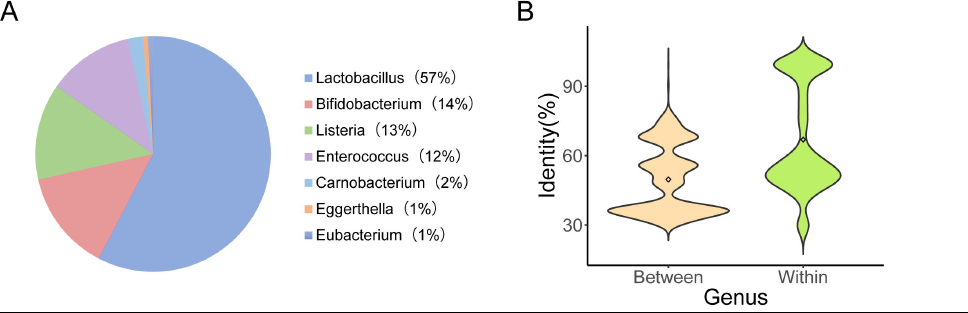
Related to Figure 2. Genera characterization of BSHs obtained from NCBI. (A) The proportion of BSHs-containing genus. (B) The identity of BSHs between genus and within genus.

**Figure S3.**
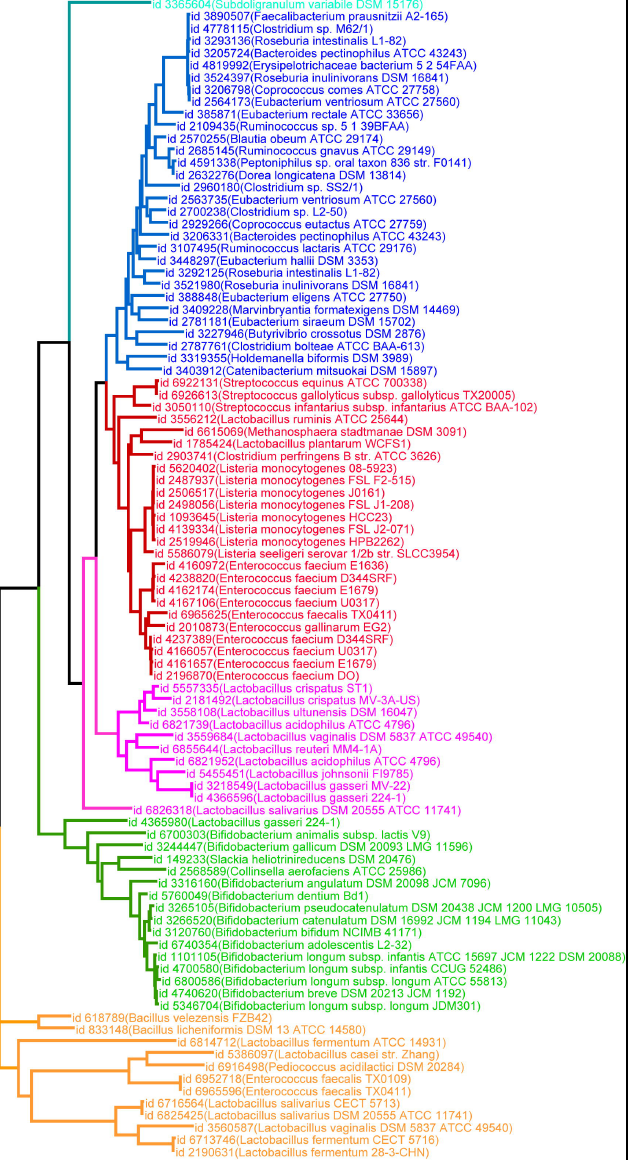
Related to Figure 4. Phylogenetic tree of BSHs in 11 populations. The id and strain name of BSHs are provided. Different color represents different phylotype of BSHs after reclassifying, from top to bottom are BSH-T01, BSH-T1, BSH-T2, BSH-T3, BSH-T4, BSH- T02.

**Figure S4.**
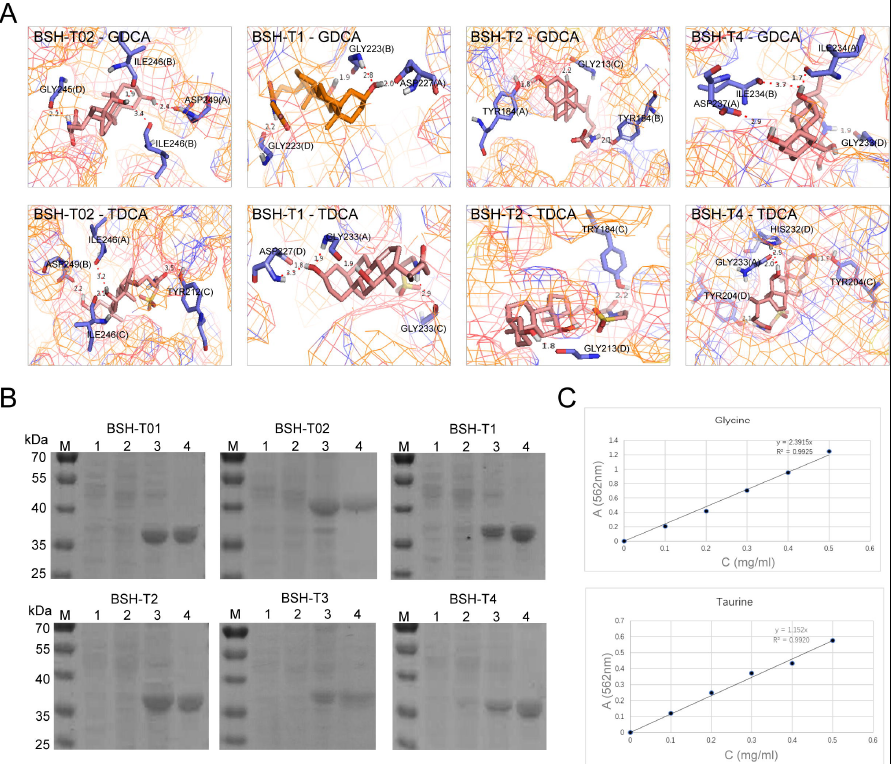
Related to Figure 5. Molecular docking and enzyme activity assay. (A) The Autodock predicted binding models of BSH-T02 to BSH-T4 with bile acids (GDCA, TDCA). (B) SDS-PAGE of BSH protein. Lane 1: BL21(DE3); Lane 2: BL21(DE3)-*bsh* (-IPTG), Lane 3: BL21(DE3)-*bsh* (+IPTG), Lane 4: BSH (purified). (C) Standard curve of glycine and taurine.

**Figure S5.**
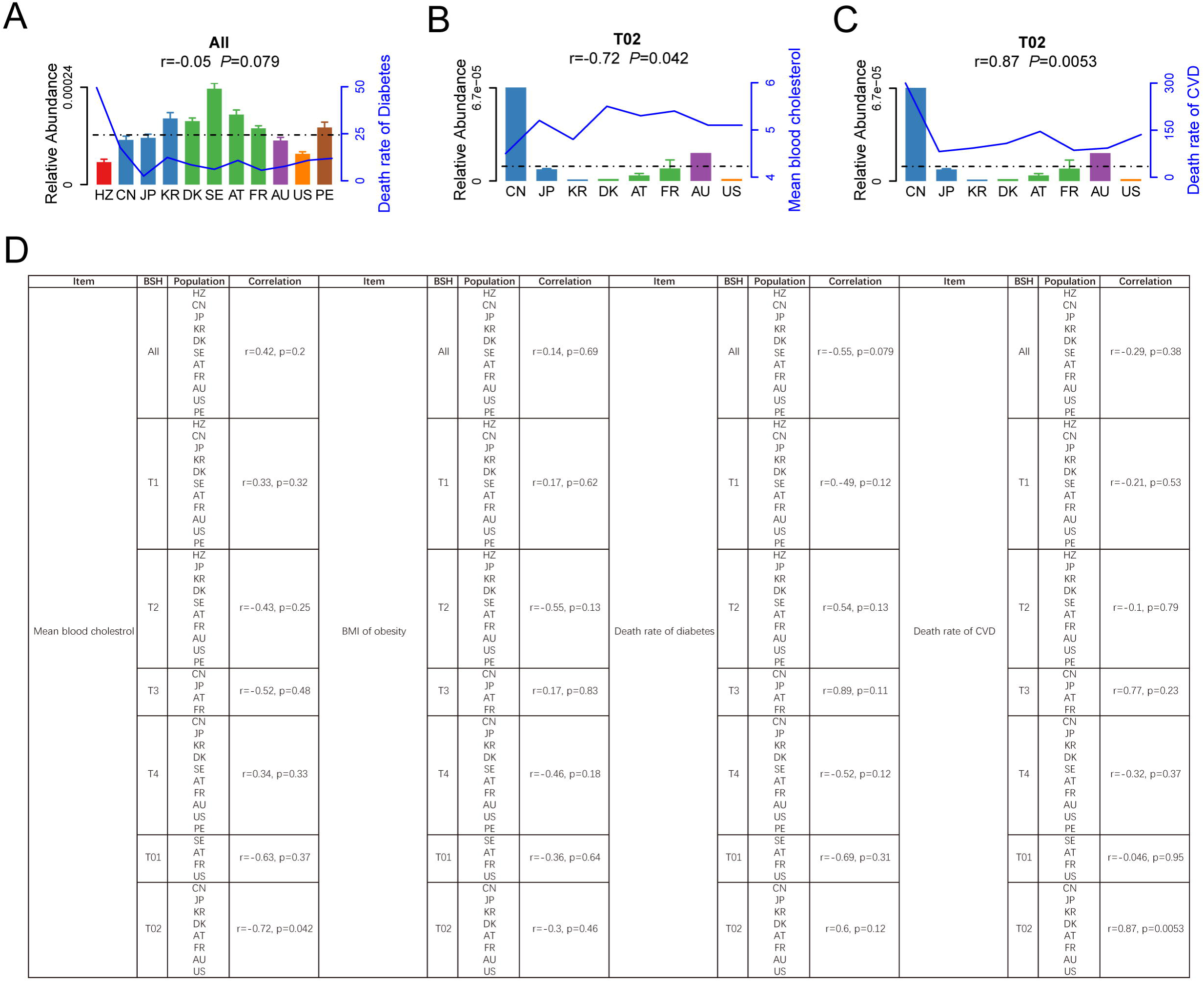
Related to Figure 4 and Figure 5. Correlation analysis of BSHs abundance and WHO released phenotypes in 11 populations. Correlation that shown significant difference between relative abundance of (A) BSH and death rate of diabetes; (B) BSH-T02 and death rate of CVD; (C) BSH-T02 and Mean blood cholesterol; (D) Correlation between different phylotypes of BSH abundance and mean blood cholesterol, BMI of obesity, death rate of diabetes and CVD.

**Table S1. Strains behave BSHs paralogs**

**Table S2. The number of individuals from the 11 countries used in this study**

**Table S3. Cumulative abundance of BSHs in different genus in 11 populations**

**Table S4. Relative abundance of 95 BSHs in 11 populations**

**Table S5. Information of 6 represent BSHs in molecular docking and enzyme activity assay**

